# Swapping of transmembrane domains in the epithelial calcium channel TRPV6

**DOI:** 10.1101/141523

**Authors:** Appu K. Singh, Kei Saotome, Alexander I. Sobolevsky

**Author notes:** Correspondence and requests for materials should be addressed to A.I.S.). These authors contributed equally to this work (Appu K. Singh, Kei Saotome).

## Abstract

Tetrameric ion channels have either swapped or non-swapped arrangements of the S1-S4 and pore domains. Here we show that mutations in the transmembrane domain of TRPV6 can result in conversion from a domain-swapped to non-swapped fold. These results reveal structural determinants of domain swapping and raise the possibility that a single ion channel subtype can fold into either arrangement *in vivo*, affecting its function in normal or disease states.

---

Transient receptor potential (TRP) channels comprise a large and functionally versatile superfamily of cation permeable ion channels that respond to a myriad of sensory modalities^1^. Structurally, TRP channels are assemblies of four identical subunits arranged symmetrically around the pore axis^2–6^. The transmembrane domain of TRP channels includes six transmembrane helices (S1-S6) and a pore loop (P-loop) between S5 and S6. The first four helices (S1-S4 domain) form a bundle that plays the role of voltage sensing domain in voltage-gated potassium channels^7^. S5, P-loop and S6 form the KcsA channel-like pore domain, with the ion conduction pathway lined by the S6 helices and extracellular loops connecting the P-loop helix to S5 and S6. In the cryo-EM reconstructions of the vanilloid subtype TRP channels TRPV1^2^ and TRPV2^3,6^, the S1-S4 domain of one subunit is packed against the pore domain of the neighboring subunit, producing a domain-swapped arrangement similar to the voltage-gated potassium channel chimera Kv1.2–2.1^7,8^.

TRPV6 and its close relative TRPV5 are TRP channels that are distinctly Ca^2+^-selective and play an important role in calcium homeostasis^9,10^. Altered expression of TRPV6 is associated with numerous human diseases, including various forms of cancer^11^. Accordingly, TRPV6 has emerged as an important target for drug design^12^. We recently solved the structure of a TRPV6 construct engineered to produce well diffracting crystals^5^. On the basis of electron density and cysteine crosslinking experiments, we built a model of TRPV6 in which the S1-S4 domain of one subunit is packed against the pore domain of the same subunit, creating a non-swapped architecture distinct from TRPV1^2^ and TRPV2^3,6^. Subsequent studies of HCN^13^, CNG^14^, Slo1^15^, Slo2.2^16^ and Eag1^17^ have revealed the prevalence of non-swapped transmembrane domain architecture in tetrameric potassium channels.

Given the domain-swapped architectures of TRPV1^2^ and TRPV2^3,6^, and relatively high sequence conservation in the transmembrane region (Supplementary Fig. 1), the absence of domain-swapping in TRPV6 was a mystery that warranted further study. We initially wondered whether the mutations present in the TRPV6 crystallization construct^5^ (TRPV6_cryst_) affected the domain arrangement. Compared to wild type, TRPV6_cryst_ has a 59-residue C-terminal truncation and four point mutations, including L495Q, which resides in the S5 transmembrane helix (Supplementary Fig. 1). This L495Q substitution, which was a spontaneous mutation that occurred during gene synthesis, was originally included in TRPV6_cryst_ because it expressed at 3-fold higher levels (Supplementary Fig. 2) than the same construct with the substitution reverted to the native leucine (TRPV6*). To examine whether the distinct non-swapped arrangement of TRPV6_cryst_ resulted from the L495Q mutation, we were compelled to solve the structure of TRPV6*.

To crystallize TRPV6*, we used the same expression system, purification protocol and crystallization conditions as for TRPV6_cryst_^5^. TRPV6* crystallized in a nearly identical crystal lattice to TRPV6_cryst_ (Supplementary Table 1) and the overall shape of the molecule was very similar (Fig. 1). Despite these similarities, to our great surprise, the structure of TRPV6* exhibited the domain swapped fold observed in other TRP channels of known structure. Attempts to use the “non-swapped” TRPV6_cryst_ model for molecular replacement resulted in massive negative and positive signals in the Fo-Fc density maps. Similar inconsistency between the crystallographic data and the model was observed when we attempted to use the “swapped” molecular replacement model of TRPV6* to solve the structure of TRPV6_cryst_. Therefore, the crystallographic data unambiguously demonstrates that the TRPV6* structure is domain-swapped. Apart from the distinct connectivity between S4 and S5 as well as S6 and TRP helix, the domain-swapped TRPV6* and non-swapped TRPV6_cryst_ structures were almost identical. Superposition of TRPV6* and TRPV6_cryst_ excluding regions of S4-S5 (residues 464–497) and S6-TRP helix (residues 577–589) gave a root mean square deviation (RMSD) of 0.865 Å.

**Figure 1.**
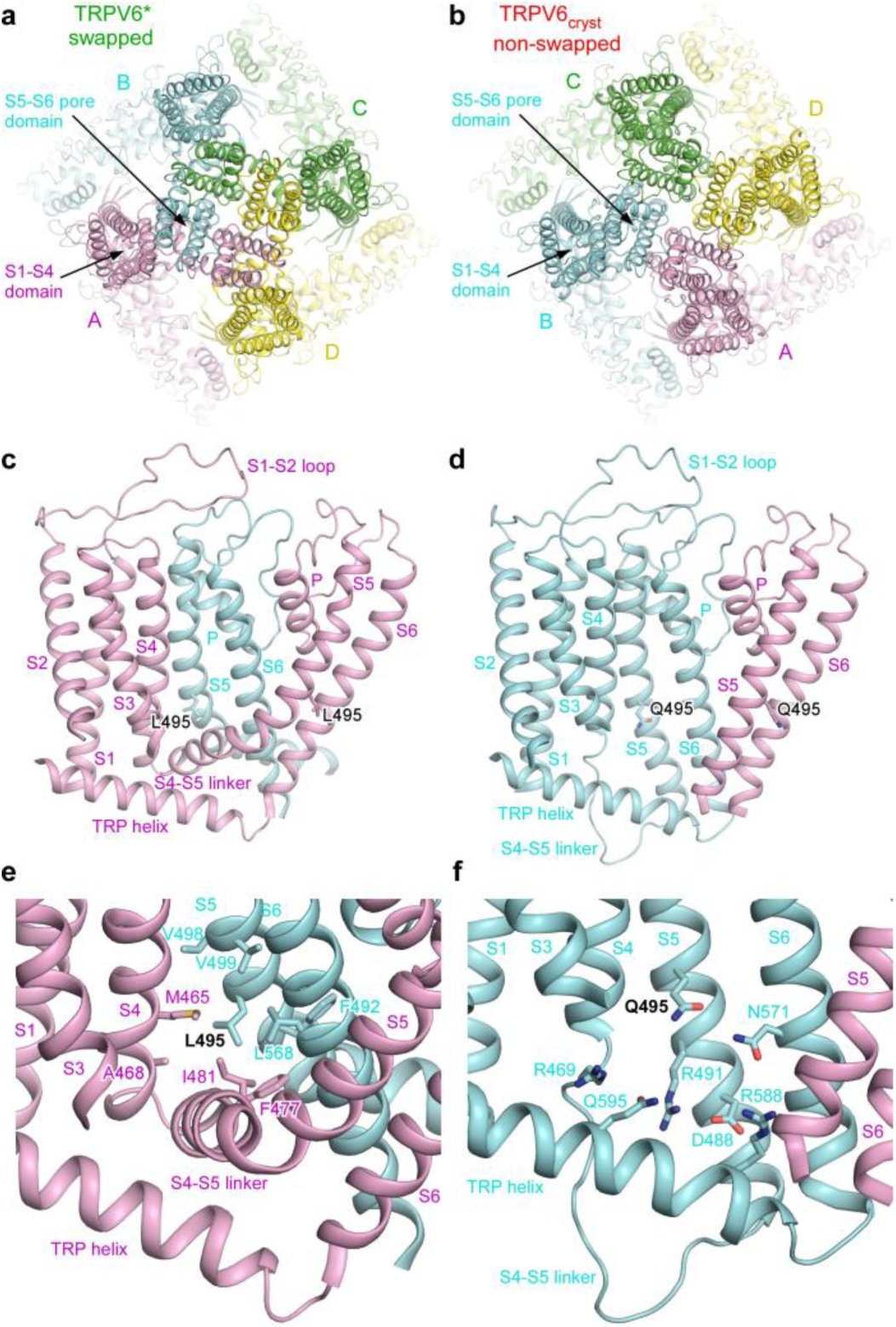
Non-swapped domain arrangement as a result of single residue substitution. **a-b**, Extracellular views of TRPV6* (**a**) and TRPV6_cryst_ (**b**), with four subunits (A-D), each shown in different color. Note, the peripheral S1-S4 domains are adjacent to the S5-S6 pore domains of the same subunit in TRPV6_cryst_, while the adjacent domains are from neighboring subunits in TRPV6*. **c-d**, Side views of one S1-S4 domain and two S5-S6 pore domains illustrating S4-S5 and S6-TRP helix connectivity unique to swapped (**c**) and non-swapped (**d**) architectures. **e-f**, Close-up views of the regions surrounding L495 in TRPV6* (**e**) and Q495 in TRPV6_cryst_ (**f**).

We reproduced crystals of TRPV6_cryst_ and confirmed its non-swapped conformation. Notably, we were able to collect new crystallographic data, where both the S4-S5 and S6-TRP helix connectivity in TRPV6_cryst_ is unambiguously resolved in the 2Fo-Fc and Fo-Fc omit maps (Supplementary Fig. 3), in stark contrast to our previous data^5^, where density for thirteen residues in S4-S5 linker (F^471^-Q^483^) was missing. Similarly, both the 2Fo-Fc and robust Fo-Fc omit maps for TRPV6* demonstrated completely different connectivity of transmembrane domains (Supplementary Fig. 3) compared to TRPV6_cryst_. These structural data therefore suggest that the single leucine L495 to glutamine substitution in S5 is responsible for the absence of domain swapping in TRPV6_cryst_. Because the sequence of TRPV6* is closer to wild-type, the domain-swapped architecture of TRPV6 likely represents the physiologically relevant domain arrangement.

The presence or absence of transmembrane domain swapping could impact the functional behavior of TRPV6 considerably. Nonetheless, ratiometric fluorescence measurements of cells loaded with the Ca^2+^ sensitive dye Fura-2 showed that the three constructs, TRPV6*, TRPV6_cryst_ and wild type TRPV6, demonstrate pore permeability to Ca^2+^ and ion channels block by Gd^3+^ (Fig. 2). Indeed, the pore architectures of TRPV6_cryst_ and TRPV6* are indistinguishable (Supplementary Fig. 4a-b), and anomalous signals for Ca^2+^ and Gd^3+^ indicated that the positions of the pore cation binding sites are similar in TRPV6* (Supplementary Fig. 4c-f) and TRPV6_cryst_^5^. Thus, the structural and functional integrity of the TRPV6 pore is overall preserved in both domain-swapped and non-swapped folds. Despite these gross similarities, kinetic analysis indicated the rate of increase in Fura-2 signal is faster in TRPV6* compared to TRPV6_cryst_ (Supplementary Fig 5). Therefore, the Ca^2+^ entry function is somewhat compromised in the non-swapped TRPV6_cryst_ construct, presumably due to differences in allosteric communication between the pore domain and the rest of the protein.

**Figure 2.**
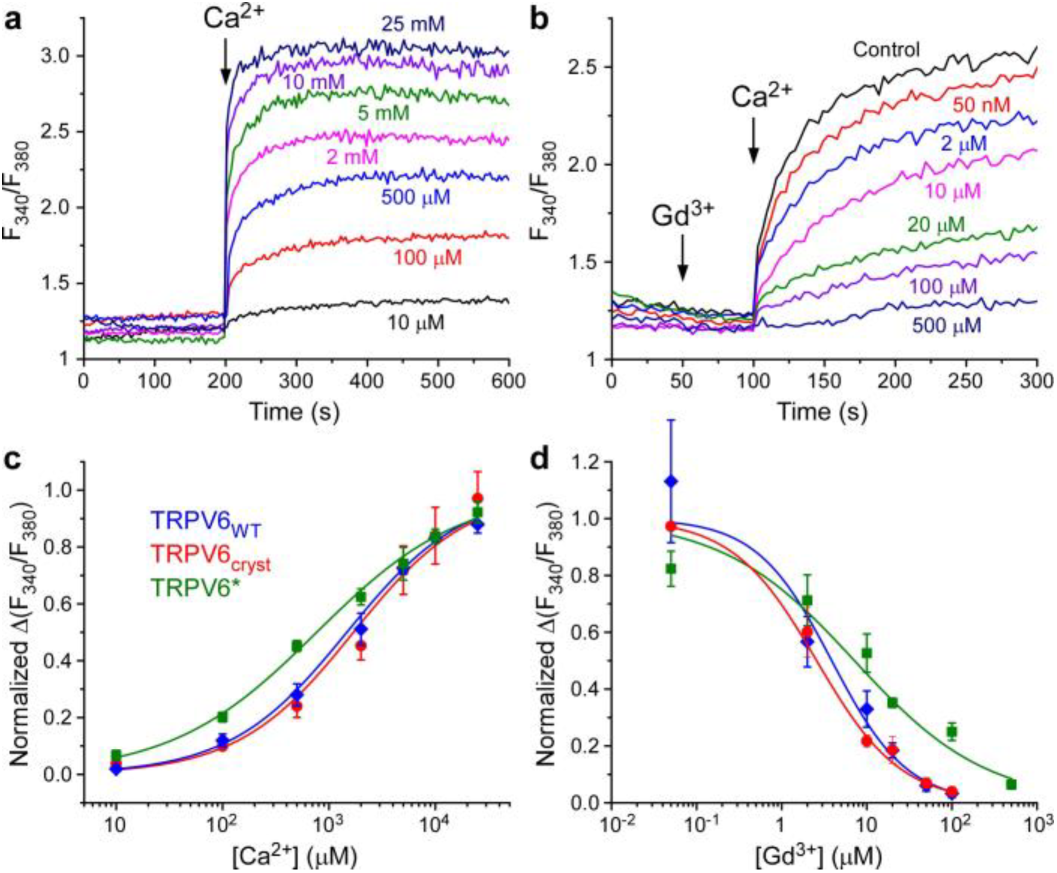
Ca^2+^ permeability and Gd^3+^ block are preserved in TRPV6 constructs with distinct domain arrangements. **a-b**, Representative ratiometric fluorescence measurements for HEK cells expressing TRPV6*. Arrows indicate the time at which the corresponding ion was added. After resuspending the cells in nominally calcium-free buffer, addition of Ca^2+^ resulted in robust concentration-dependent increase in Fura-2 signal (**a**). In contrast, pre-incubation of cells with increasing concentrations of Gd^3+^ resulted in concentration-dependent reduction in Fura-2 signal (**b**). **c-d**, Normalized changes in F340/F380 at varying concentrations of Ca^2+^ (**c**) and Gd^3+^ (**d**) calculated for wild type TRPV6 (blue), TRPV6_cryst_ (red) and TRPV6* (green). In **d**, Δ F340/F380 was measured after preincubation with Gd^3+^ and subsequent addition of 2 mM Ca^2+^.

How does the L495Q mutation result in the non-domain swapped fold of TRPV6? In TRPV6*, L495 projects its side chain into a hydrophobic pocket formed by residues of S5 and S6 of one subunit and residues at the S4-S5 elbow of the adjacent subunit (Fig. 1e). By contrast, in TRPV6_cryst_, Q495 is rotated by ~120° with respect to the orientation of L495 in TRPV6* and forms polar interactions with the hydrophilic pocket formed by S4-S6 and the TRP helix of the same subunit (Fig. 1f). These observations suggest that hydrophobic burial of the L495 residue plays an important role in driving the domain-swapped fold of TRPV6. Disruption of these hydrophobic interactions leads to the non-swapped fold. In other members of TRPV family, either leucine or valine occupy the position homologous to L495 in TRPV6 (Supplementary Fig. 1b). Further, a hydrophobic pocket similar to TRPV6 is also present in the structures of TRPV1/2^2,3,6^, suggesting a conserved role for this region in determining the domain arrangement in TRPV channels.

In the case of non-swapped potassium channels, the absence of domain-swapping was attributed to shorter S4-S5 linkers^13–17^. We tested whether TRPV6 domain arrangement could be biased toward a non-swapped conformation by shortening of the S4-S5 linker, which has a well-defined helical structure in the domain-swapped TRPV6* but lacks secondary structure in TRPV6_cryst_. Remarkably, TRPV6* constructs with up to ten residues deleted in the S4-S5 linker expressed robustly and could be isolated as a monodisperse species with gel filtration elution time corresponding to the tetrameric assembly (Supplementary Fig. 6a-b). We determined the crystal structure of TRPV6* with four residues deleted in the S4-S5 linker (TRPV6*-del1) and confirmed its non-swapped architecture (Supplementary Fig. 6c-d). Therefore, the length of the S4-S5 linker is indeed a critical determinant of domain-swapping. Interestingly, ratiometric fluorescence measurements did not detect measurable Ca^2+^ influx in HEK cells expressing TRPV6*-del1, suggesting that while the tetrameric expression and assembly in this mutant is preserved, gating function is significantly perturbed.

In summary, our studies demonstrate that small sequence changes can result in a major alteration of the transmembrane domain arrangement of a tetrameric ion channel. Depending on the nature of the mutation, this structural alteration can lead to strong or weak changes in ion channel function. We speculate that a single tetrameric ion channel subtype can adopt swapped or non-swapped transmembrane domain arrangements *in vivo*, depending on the presence of mutations, post-translational modifications, or other perturbations. The impact of changes in domain-swapping on ion channel function may play a role in pathophysiology.

## Methods

### Constructs

Our previous crystallization construct TRPV6_cryst_^5^ comprised of residues 1–668 of rat TRPV6 and contained the point mutations I62Y, L92N, M96Q and L495Q. We made a new construct TRPV6*, which is identical to TRPV6_cryst_ except the reversal of the single L495Q mutation in S5 to the native leucine residue. To understand the role of S4-S5 linker in domain swapping, TRPV6*-del1, TRPV6*-del2, TRPV6*-del3 and TRPV6*-del4 constructs (Supplementary Fig. 6a) were prepared by deleting four (477–480), six (475–480) eight (473–480) and ten (471–480) residues in S4-S5 linker, which were disordered in the original TRPV6_cryst_ structure^5^.

### Expression and Purification

TRPV6 constructs described in this paper were expressed and purified as previously described for the TRPV6_cryst_^5^. In brief, TRPV6 constructs were introduced into the pEG BacMam vector with C-terminal thrombin cleavage site (LVPRG) followed by eGFP and streptavidin affinity tag (WSHPQFEK), for the expression in suspension of baculovirus-transduced HEK293 GnTI^−^ cells. 48–72 hours after transduction with P2 virus, cells were harvested, washed with PBS at pH 8.0 and after sonication, cellular membranes were prepared. The protein was extracted from cellular membranes using 20 mM n-dodecyl-β-D-maltopyranoside, and purified using Strep-Tactin affinity chromatography followed by size exclusion chromatography.

### Crystallization and structure determination

Optimized crystals of purified TRPV6* or TRPV6*-del1 were grown in the same condition as crystals of TRPV6_cryst_^5^, which included a reservoir solution consisting of 20–24% PEG 350 MME, 50 mM NaCl and 50 mM Tris-HCl pH 8.0–8.5 in hanging drop configuration at 20°C. Prior to crystallization using hanging drop method, the protein was subjected to ultracentrifugation (Ti100 rotor, 40000 rpm, 40 min, 4°C) to remove the aggregated protein. Crystals were cryoprotected by serial transfer into the buffer composed of 100 mM NaCl, 100 mM Tris-HCl pH 8.2, 0.5 mM DDM and 50 mM ammonium formate, containing increasing concentration of PEG 350 MME, with maximum concentration of up to 33–36%, and then flash frozen in liquid nitrogen. To obtain crystals with Ca^2+^ or Gd^3+^, protein was incubated with 10 mM CaCl2 or 1 mM GdCl3, respectively, for at least 1 hour at 4°C prior to crystallization.

X-ray diffraction data collected at APS (beamlines 24-ID-C/E), NSLSII (beamline 17-ID) or ALS (beamline 5.0.2) were indexed, integrated and scaled using XDS^18^ or HKL2000^19^. The initial phase information and structures were solved by molecular replacement using Phaser^20^ and the structure of rat TRPV6_cryst_^5^ (PDB ID: 5IWK) as a search probe. Most of TRPV6*, including the ankyrin domain, S1-S4, pore module and C-terminal hook, was nearly identical to the TRPV6_cryst_; the rest of the structure was built using the omit map as a guide. The robust electron density for the S4-S5 linker was evident from the initial phases obtained by molecular replacement, and map features improved further during refinement. The model was refined by alternating cycles of building in COOT^21^ and automatic refinement in Phenix^22^ and Refmac^23^. All structural figures were made using PyMOL^23^ (www.pymol.org)

### Fura 2-AM measurements

The intracellular Ca^2+^ measurements from HEK cells expressing rat TRPV6 or TRPV6_cryst_ or TRPV6* were performed as described previously^5^. All Fura2-AM-based fluorescence measurements were conducted using spectrofluorometer QuantaMaster™ 40 (Photon Technology International) at room temperature in a quartz cuvette under constant stirring.

---

**Supplementary Figure 1.**
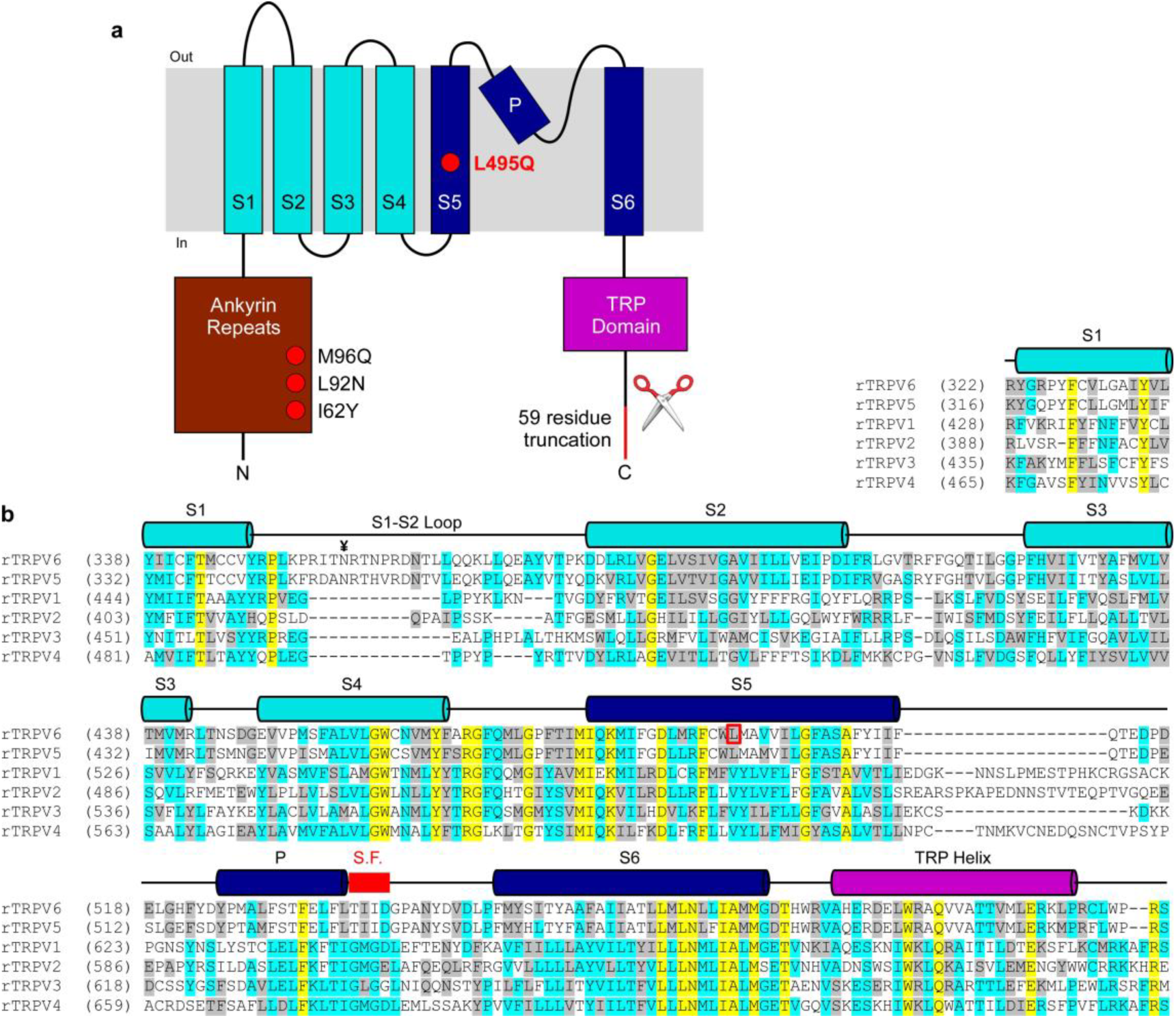
Topology and sequence alignment of TRPV subunits. **a**, Membrane topology of TRPV subunits. Red circles and red line indicate four point mutations and C-terminal deletion in TRPV6_cryst_ construct, respectively. **b**, Sequence alignment of the transmembrane domain regions in rat TRPV subtypes. Helices are indicated by cylinders above the sequence. L495 (open red box) and the selectivity filter in TRPV6 (thick red line) are highlighted. ¥ marks the N-linked glycosylation site in the extracellular loop connecting S1 and S2 conserved in TRPV6 (and TRPV5) channels.

**Supplementary Figure 2.**
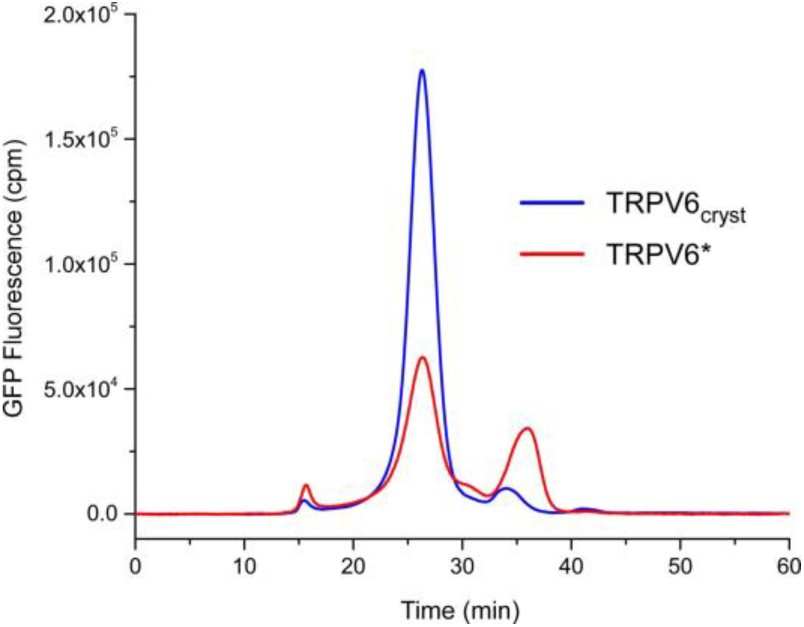
FSEC traces for crudely solubilized TRPV6_cryst_ and TRPV6*. FSEC traces were recorded using Superose 6 size-exclusion chromatography column monitoring GFP fluorescence. Note the difference in the peak height, indicating approximately 3 fold greater expression levels of TRPV6_cryst_ compared to TRPV6*.

**Supplementary Figure 3.**
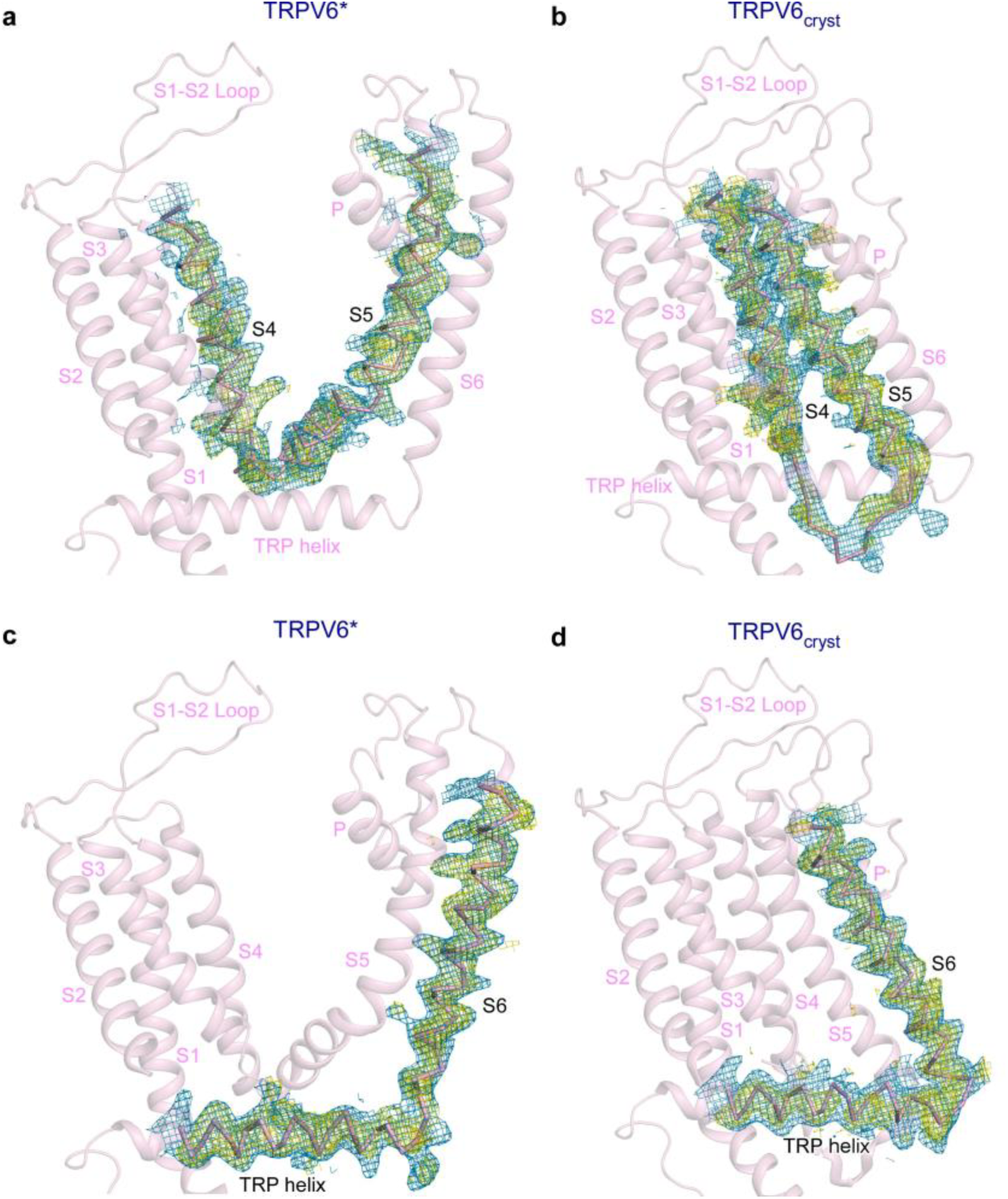
Dramatic change in TRPV6 transmembrane domain connectivity as a result of single-residue substitution in S5. **a-d**, Transmembrane portions of TRPV6* (leucine at position 495, left) and TRPV6_cryst_ (glutamine at position 495, right) viewed parallel to the membrane. S4-S5 (upper row) and S6-TRP helix (bottom row) connectivity is illustrated by 2Fo-Fc (blue mesh, 1σ) and omit Fo-Fc (yellow mesh, 2.5σ) maps.

**Supplementary Figure 4.**
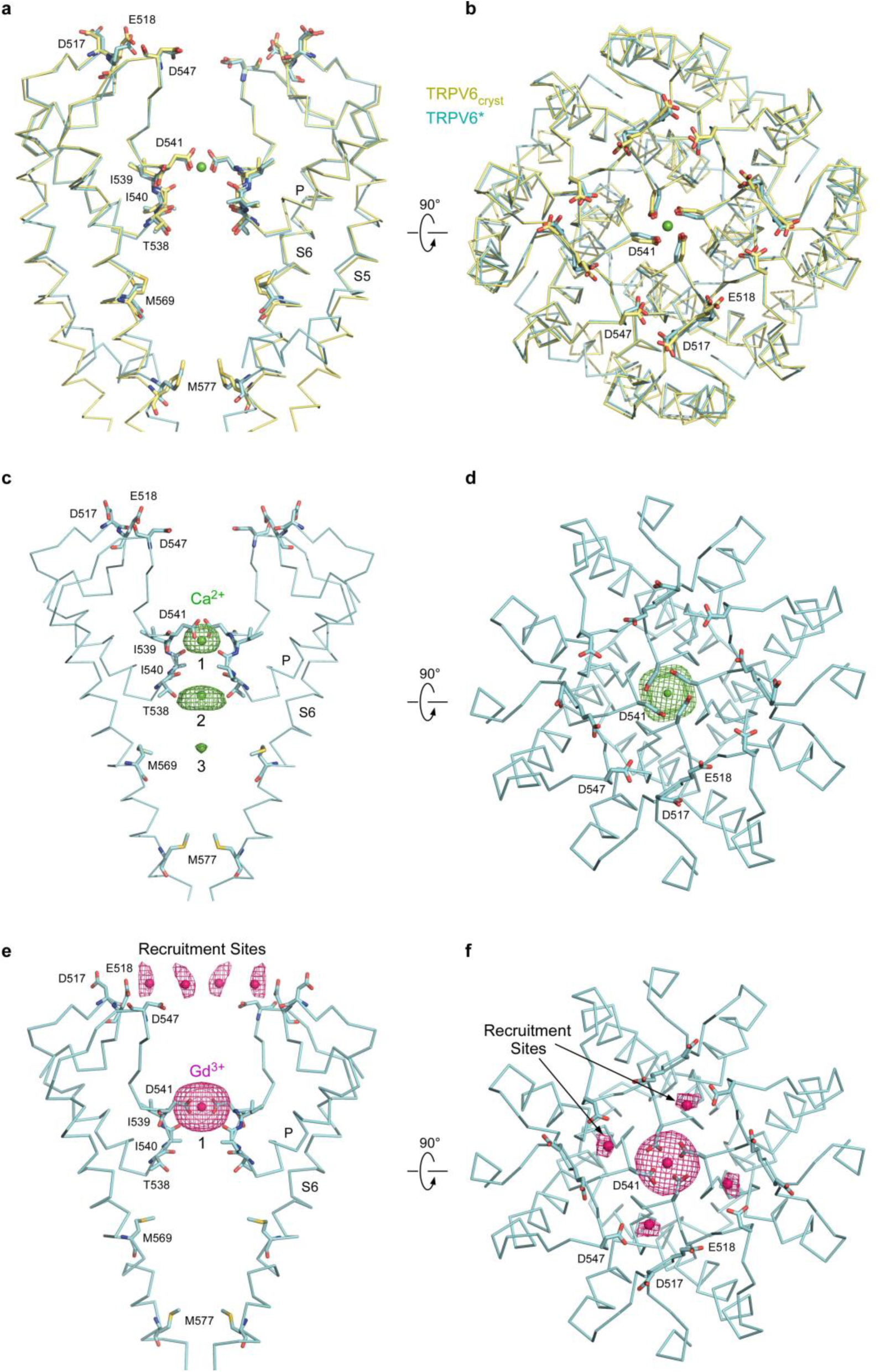
TRPV6 structures with swapped and non-swapped transmembrane domains have identical ion channel pores. **a-b**, Superposition of the pore domains in TRPV6* (cyan) and TRPV6_cryst_ (yellow) viewed parallel to the membrane (**a**) or extracellularly (**b**), with residues important for cation binding shown in stick representation. Calcium atom is shown as a green sphere. **c-f**, Side (**c,e**) and top (**d,f**) views of the TRPV6* pore. Front and back subunits in c and e are removed for clarity. Green and pink mesh shows anomalous difference electron density for Ca^2+^ (**c-d**, 3σ) and Gd^3+^ (**e-f**, 3.5σ), respectively, and ions are shown as spheres of the corresponding color. Positions of cation binding sites are similar to TRPV6_cryst_^5^.

**Supplementary Figure 5.**
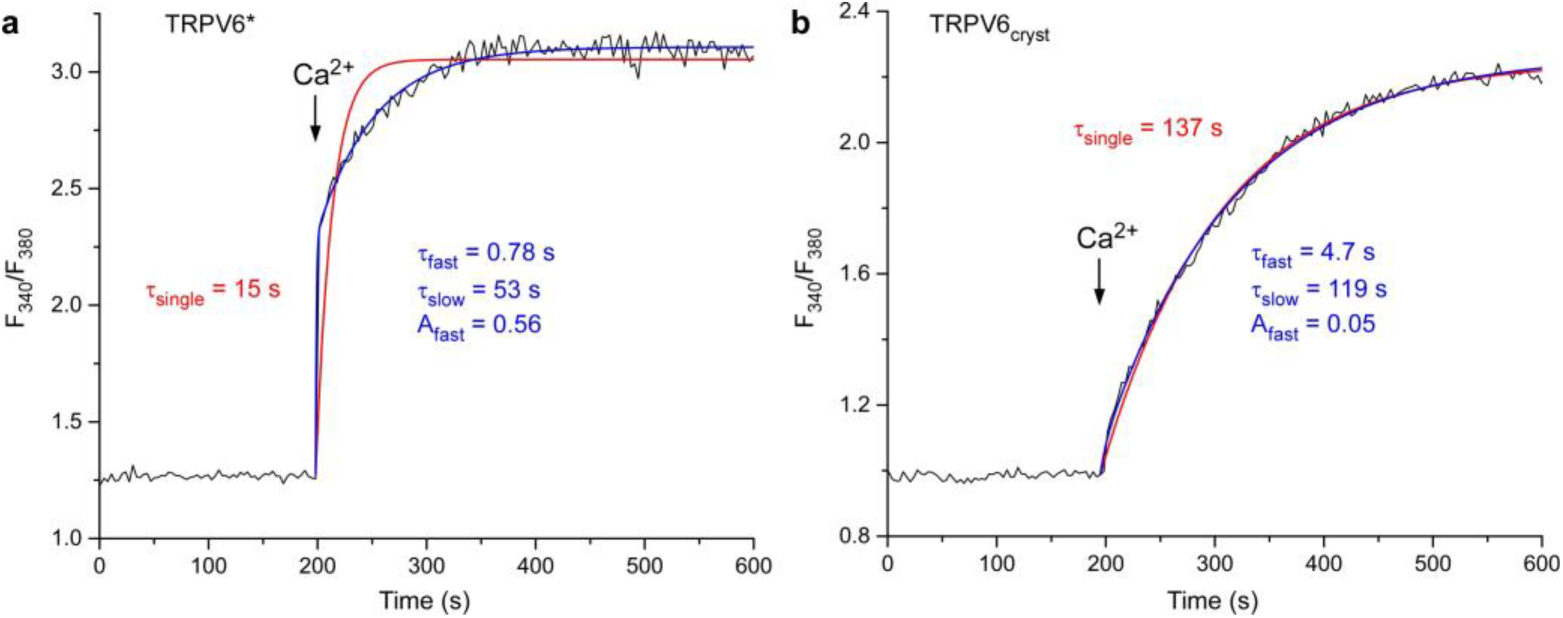
Different kinetics of ratiometric curves for TRPV6* or TRPV6_cryst_ at physiological extracellular calcium concentration. **a-b**, Representative ratiometric fluorescence measurements for HEK cells expressing TRPV6* (**a**) or TRPV6_cryst_ (**b**). Arrows indicate the time at which 2 mM Ca^2+^ was added. The curves were fitted with one (red) or two (blue) exponentials. Note, TRPV6* kinetics was well described by the double but not the single exponential. Over three measurements, the time constants of the single exponential fit (τ_single_) were 14.5 ± 2.7 s for TRPV6* and 110 ± 16 s for TRPV6_cryst_, τ_fast_ = 2.2 ± 1.0 s, τ_slow_ = 54 ± 11 s and A_fast_ = 0.57 ± 0.01 for TRPV6* and τ_fast_ = 5.4 ± 2.8 s, τ_slow_ = 135 ± 23 s and A_fast_ = 0.10 ± 0.03 for TRPV6_cryst_. At n = 3, the values of τ_single_ and A_fast_ for TRPV6* and TRPV6_cryst_ were statistically different (t-test, P < 0.05).

**Supplementary Figure 6.**
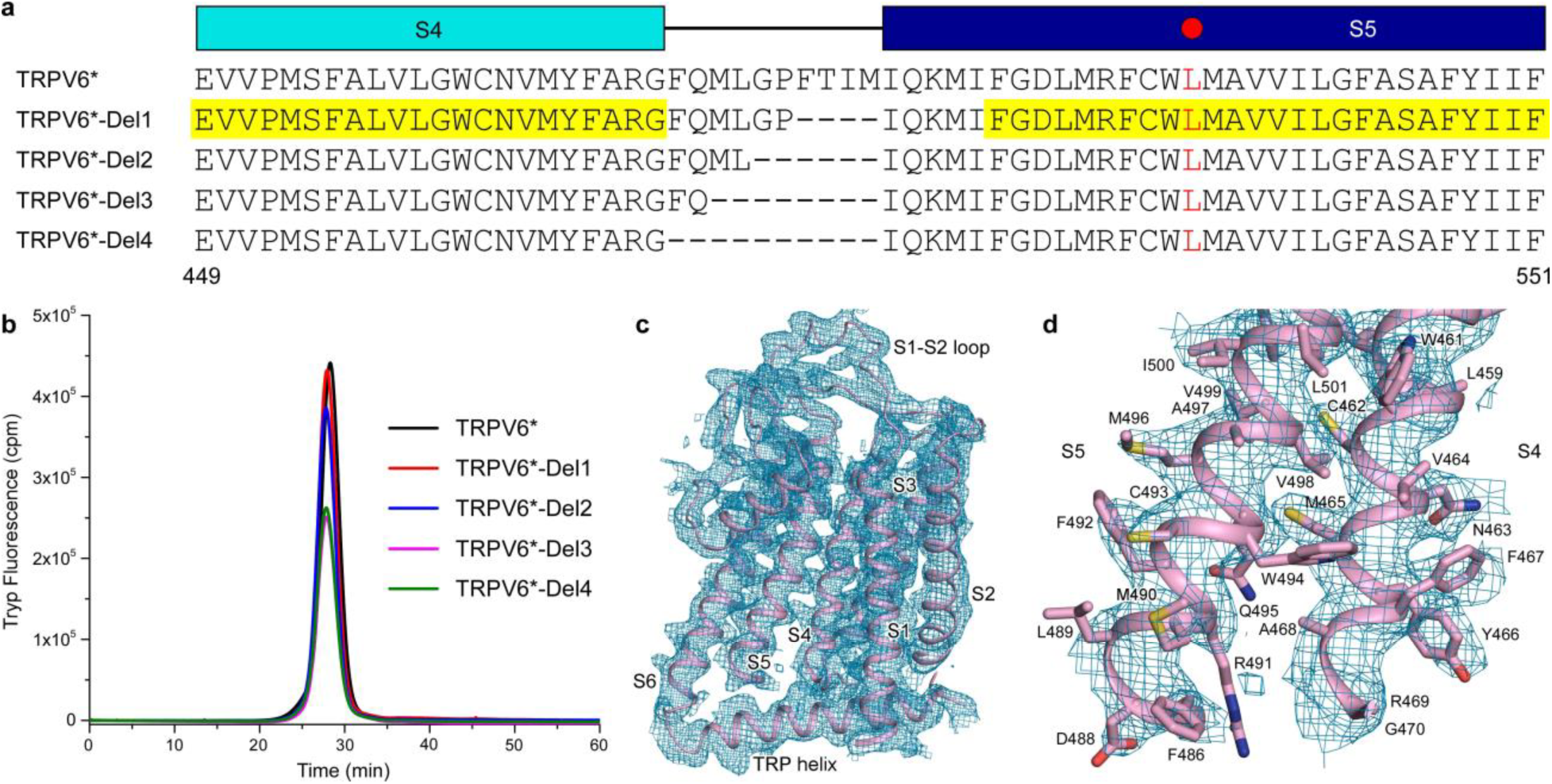
Deletions in S4-S5 preserve TRPV6 tetrameric assembly and generate non-swapped topology. **a**, Sequence alignment of TRPV6* and mutants with deletions in the S4-S5 linker. Leucine L495 is indicated by the red circle. The sequence corresponding to the resolved regions in TRPV6*-Del1 structure is highlighted by yellow. **b**, FSEC traces for purified TRPV6* without and with deletions in S4-S5 following tryptophan fluorescence. Note, all samples demonstrate a single monodisperse tetrameric peak. **c**, Transmembrane domain of a single TRPV6*-Del1 subunit demonstrates a non-swapped architecture with the characteristic S6-TRP helix connectivity. Blue mesh shows 2Fo-Fc map (1σ). **d**, Close-up view of the S4-S5 illustrating boundaries of a missing region (residues 471–485).

**Supplementary Table 1.** Data collection and refinement statistics.

|  | TRPV6* | TRPV6<sub>cryst</sub> | TRPV6*-Del1 | TRPV6*-Ca<sup>2+</sup> | TRPV6*-Gd<sup>3+</sup> |
| --- | --- | --- | --- | --- | --- |
| **Data Collection** |  |  |  |  |  |
| Beamline | APS-24ID-C | APS-24ID-C | APS-24ID-C | APS-24ID-C | APS-24ID-C |
| Space group | P42<sub>1</sub>2 | P42<sub>1</sub>2 | P42<sub>1</sub>2 | P42<sub>1</sub>2 | P42<sub>1</sub>2 |
| Cell dimensions | 146.03 | 145.53 | 145.32 | 145.48 | 144.02 |
| a, b, c, (Å) | 146.03 | 145.53 | 145.32 | 145.48 | 144.02 |
|  | 116.05 | 114.13 | 113.67 | 110.50 | 114.34 |
| $\alpha, \beta, \gamma$ (°) | 90 90 90 | 90 90 90 | 90 90 90 | 90 90 90 | 90 90 90 |
| Wavelength (Å) | 0.9791 | 0.9791 | 0.9791 | 1.75 | 1.00 |
| Resolution (Å)* | 50.0 - 3.25<br>(3.44 - 3.25) | 50.00 - 3.45<br>(3.56 - 3.45) | 50.0 - 3.40<br>(3.52- 3.40) | 50.0 – 3.6<br>(3.68 - 3.60) | 50.0 - 3.85<br>(3.99 - 3.85) |
| Completeness<br>(%)* | 98.4<br>(97.2) | 99.8<br>(99.6) | 99.26<br>(95.13) | 99.5<br>(95.1) | 95.9<br>(91.8) |
| Redundancy* | 8.6<br>(7.1) | 8.5<br>(8.5) | 8.5<br>(6.7) | 18.9<br>(17.3) | 8.3<br>(5.5) |
| $\ I/\sigma\ $ * | 18.31<br>(1.05) | 11.3<br>(1.1) | 11.44<br>(1.22) | 11.9<br>(1.07) | 12.1<br>(1.10) |
| R<sub>meas</sub> (%)** | 8.7<br>(190.9) | 9.93<br>(194.45) | 13.0<br>(140.9) | 14.6<br>(260.5) | 19.4<br>(255.8) |
| CC<sub>1/2</sub> | 99.8<br>(60.3) | 99.6<br>(44.65) | 99.9<br>(61.8) | 99.6<br>(67.1) | 99.5<br>(42.9) |
| **Refinement** |  |  |  |  |  |
| Resolution (Å)* | 50.0 - 3.25<br>(3.36 - 3.25) | 49.91 - 3.45<br>(3.66 - 3.45) | 50.00 - 3.40<br>(3.52-3.40) | 50.0 – 3.6<br>(3.68 - 3.60) | 50.0 - 3.85<br>(3.99 - 3.85) |
| Completeness<br>(%) | 96<br>(93.8) | 99.8<br>(98.5) | 99.26<br>(95.13) | 96<br>(93.8) | 97.86<br>(99.20) |
| Number of<br>reflections | 18531<br>(1724) | 17758<br>(791) | 17211<br>(1605) | 18531<br>(1724) | 20034<br>(1973) |
| R<sub>work</sub>/R<sub>free</sub> | 0.273/0.297 | 0.278/0.298 | 0.286/0.305 | 0.273/0.297 | 0.275/0.295 |
| Number of atoms |  |  |  |  |  |
| Total | 4722 | 4809 | 4727 | 4722 | 4897 |
| Ligand | 16 | 3 | 16 | 16 | 155 |
| B-factor (Å<sup>2</sup>) |  |  |  |  |  |
| Protein | 120.5 | 147.07 | 145.47 | 120.5 | 150.11 |
| Ligand | 77.27 | 86.12 | 164.66 | 77.27 | 164.71 |
| **RMS deviations** |  |  |  |  |  |
| Bond length (Å) | 0.003 | 0.003 | 0.009 | 0.003 | 0.002 |
| Bond angles (°) | 0.7 | 0.84 | 0.84 | 0.7 | 0.51 |
| **Ramachandran** |  |  |  |  |  |
| Favored (%) | 94.6 | 93.6 | 94.6 | 94.6 | 94.6 |
| Allowed (%) | 5.4 | 5.9 | 4.9 | 5.4 | 4.8 |
| Disallowed (%) | 0.0 | 0.5 | 0.5 | 0.0 | 0.6 |

\*Highest resolution shell in parentheses. 5% of reflections were used for calculation of R<sub>free</sub>.

